# Male sex and age biases viral burden, viral shedding, and type 1 and 2 interferon responses during SARS-CoV-2 infection in ferrets

**DOI:** 10.1101/2021.01.12.426381

**Authors:** Magen E. Francis, Brian Richardson, Mara McNeil, Melissa Rioux, Mary K. Foley, Anni Ge, Roger D. Pechous, Jason Kindrachuk, Cheryl M. Cameron, Christopher Richardson, Jocelyne Lew, Mark J. Cameron, Volker Gerdts, Darryl Falzarano, Alyson A. Kelvin

## Abstract

SARS-CoV-2 (Severe Acute Respiratory Syndrome Coronavirus 2) hospitalizations and deaths disportionally affect males and the elderly. Here we investigated the impact of male sex and age by infecting adult male, aged male, and adult female ferrets with SARS-CoV-2. Aged male ferrets had a decrease in temperature which was accompanied by prolonged viral replication with increased pathology in the upper respiratory tract after infection. Transcriptome analysis of the nasal turbinates and lungs indicated that female ferrets had significant increases in interferon response genes (OASL, MX1, ISG15, etc.) on day 2 post infection which was delayed in aged males. In addition, genes associated with taste and smell such as RTP1, CHGA, and CHGA1 at later time points were upregulated in males but not in females. These results provide insight into COVID-19 and suggests that older males may play a role in viral transmission due to decreased antiviral responses.

## Introduction

In December 2019, Severe Acute Respiratory Syndrome Coronavirus 2 (SARS-CoV-2) emerged in Wuhan, Hubei, China, which has led to a global pandemic ^1^. SARS-CoV-2, identified as the causative agent of COVID-19 (Coronavirus disease 2019), is a beta-coronavirus belonging to the *Coronaviridae* family of enveloped positive sense, single-stranded RNA viruses with significant similarities to SARS-CoV which emerged in 2002 ^2,3^. Infection in humans with SARS-CoV-2 can lead to severe disease, hospitalization, and death in some cases while others may remain subclinical or develop only mild disease; therefore, the clinical picture of COVID-19 is significantly varied and disease can range from mild nasal congestion and sore throat to severe pneumonia with diffuse alveolar damage leading to multi-organ failure and death ^4–6^. Non-respiratory symptoms such as loss of taste and smell, the development of microblood clots (COVID-toes), strokes, neurological impairment, and long-term complications (COVID long-haulers) have also been reported or associated with infection ^7,8^. These symptoms are indicative of direct or indirect effects of the virus on major organ systems such as the neurosensory system and the cardiovascular system. As of November 18, 2020, there have been more than 56 million confirmed cases and over 1.3 million deaths world-wide ^9^, highlighting the urgent need for vaccines, antivirals, and a detailed understanding of the mechanisms underlying disease progression and viral transmission.

Epidemiological analysis across several countries has indicated that there are several host factors including sex, age, and co-morbidities (chronic obstructive pulmonary disease, diabetes, hypertension, and cancer) that can increase COVID-19 severity. Demographic analysis across several countries including China, France, Germany, Iran, Italy, and the US have inidicated that men have increased severe disease and mortality following SARS-CoV-2 infection compared to women ^10,11^. Age also has a clear impact on COVID-19 where case fatality rates (CFR) significantly increases with age ^4,11^. Indeed, the CFR for COVID-19 in individuals < 40 years of age is less than 0.2% whereas those aged between 60-69, 70-79, and 80+ have CFRs of 3.6%, 8.0%, and +14%, respectively ^11,12^. Thus people >65 years of age are considered the highest risk group for developing severe illness by the US Centers for Disease Control and Prevention (CDC) ^13^. In the US alone, 80% of COVID-19-related fatalities have occurred in patients >65 years of age ^4^ and long-term care facilities have been demonstrated to be particularly vulnerable to COVID-19 ^14–16^. Clinical identification of COVID-19 in the elderly is also constrained by differences in disease symptoms including tachypnea, unexplained tachycardia and increased blood pressure, altered mental status, muscle pain, and fatigue ^14,17^. Understanding COVID-19 disease mechanisms in higher-risk groups, including males and the elderly, remains a significant public health priority and will inform supportive care and treatment modalities as well as vaccine strategies for high-risk groups.

Preclinical models are essential for advancing countermeasures for infectious diseases into human clinical trials. Animal models are indispensable for responses to newly emerging infectious diseases given their importance for preclinical evaluations of vaccines and therapeutics and corresponding insights into disease pathophysiology^18^. However, development of these models requires identification of species that are susceptible to infection as well as recapitulation of human clinical disease. While mice have been traditionally employed in preclinical investigations due to reagent availability and cost, they are not typically susceptible to human clinical viral isolates without adaptation. In contrast, ferrets are commonly susceptible to human viruses, including ebolaviruses, influenza viruses and coronaviruses, and have been used for virus characterization and vaccine testing ^18–20^. Additionally, the respiratory tract of ferrets has human-like physiology including similarities in the numbers of terminal branches in the lower respiratory tract, distribution of cellular receptors for viruses, and body:lung surface area ratio ^21^. Ferrets display similar symptoms and clinical features to humans during respiratory infections, including fever, nasal discharge, coughing, and weight loss ^19,22–25^. They have also been used to investigate age-related host responses, disease severity, and pathogenic immune mechanisms during respiratory virus infection ^23,24^. Further, the utility of ferrets for infection studies has been demonstrated in SARS-CoV investigations for vaccine development and innate immune responses, including interferon stimulated genes ^19,22,26–31^.

The emergence and rapid global spread of SARS-CoV-2 has required an unparalleled search for animal models that meet the criteria needed for timely preclinical characterization of viral pathogenesis and therapeutics identification ^32^. Currently, ferrets, domesticated cats, nonhuman primates, and Syrian hamsters have been shown to be susceptible to infection and support viral replication ^33–38^. Although not naturally susceptible, genetically modified mice as well as mouse adapted SARS-CoV-2 viruses in wild type mice have also been used for SARS-CoV-2 preclinical modeling ^39–41^. Ferret infection studies have shown that the virus can be readily transmitted between animals by contact as well as aerosol transmission ^34,42^. Ferrets are sexually dimorphic animals that display sex-biased health conditions suggesting their utility for COVID-19 sex biases ^21^. Moreover, age-related viral disease severity has been shown in ferrets by our group and others ^23,24,43–46^. As COVID-19 severity has been associated with the male sex and age, we investigated these biological variables and their contribution to disease in ferrets by identifying the molecular mechanisms underlying these contributions. We characterized sex- and age-related viral load and shedding as well as immune gene expression in SARS-CoV-2 infected ferrets. Taken together, our data demonstrated that older males had a longer duration of viral shedding from the upper respiratory tract with a delayed induction of antiviral and interferon stimulated genes compared to younger females and males. These findings may suggest that antiviral treatment may be more beneficial to male patients. In addition, our findings highlight that not only sociological and societal factors, but also biological factors including sex hormones, sex chromosomes, and aging, contribute to COVID-19 sex-biases.

## Results

### SARS-CoV-2 infection causes hypothermia in male ferrets but fever in females

Significant sex and age biases have been observed for severe COVID-19 ^47–50^. Understanding the molecular mechanisms contributing to these biases is essential for therapeutic development and preventative measures which will provide a greater protection for vulnerable populations such as the elderly and males. Ferrets are often used as a preclinical model for dissecting the molecular mechanisms of human viral diseases including those with sex and age biases ^43,51^. Here, we employed ferrets to gain a better understanding of human SARS-CoV-2 infection and the increased risk of severe COVID-19 in aged males. Ferrets are typically purchased for scientific studies after they have been fixed (i.e., spayed or neutered). To address our questions surrounding the roles of sex and age in severe COVID-19 disease, we acquired intact female and male (adult and aged) ferrets from Triple F farms for our studies. Three experimental groups were employed to assess the roles of sex and age: group 1 – adult female ferrets (1 year of age); group 2 – adult male ferrets (1 year of age); and group 3 – aged male ferrets (over 2 years of age). All three groups were intranasally inoculated with 10^6^ TCID_50_ of SARS-CoV-2 while anesthetized. Clinical responses such as weight loss, temperature, activity, and other signs of neurological and gastrointestinal disease were monitored for 14 - 21 days post inoculation (pi). In-life nasal washes, blood samples and tissues were collected on day 2, 5, 7, 14, and 21 pi to analyse host responses and viral burden among the groups.

Ferrets display human-like clinical symptoms after viral infection ^52^. Weight and temperature were monitored after inoculation for 14 days (Fig. 1). Minimal clinical changes in weight were observed for all groups throughout the time course. All groups remained close to their original weight (100%) and no statistical differences were noted (Fig. 1A). Interestingly, significant variability was observed for temperature between group comparisons. Female ferrets experienced significant biphasic temperature increases after inoculation compared to their baseline values. On day 1 pi, female temperature increased to 101% of original temperature that was sustained until day 4. The temperature of female ferrets then again increased on day 9 when temperature was recorded at 102% of original readings. Although this temperature increase was mild in the females, it differed significantly from both male ferret groups. The male ferrets experienced a temperature decrease after inoculation (98-99% of original temperature). Aged male ferrets (represented by the grey line on the graph) had lower temperatures than the adult males for longer durations. Aged males exhibited a biphasic trend of initial decrease days 1-3 pi and a second decrease from days 9-12 with significantly lower temperatures than female temperatures on days 1, 2, 3, 7, 9, 10, 11, and 12 (Fig. 1B). No statistical differences were noted between the adult males and aged males although adult males had fewer statistically hypothermic days compared to females. Together, this data suggests a sex effect on temperature change exacerbated by increases in age following SARS-CoV-2 infection.

**Fig. 1:**
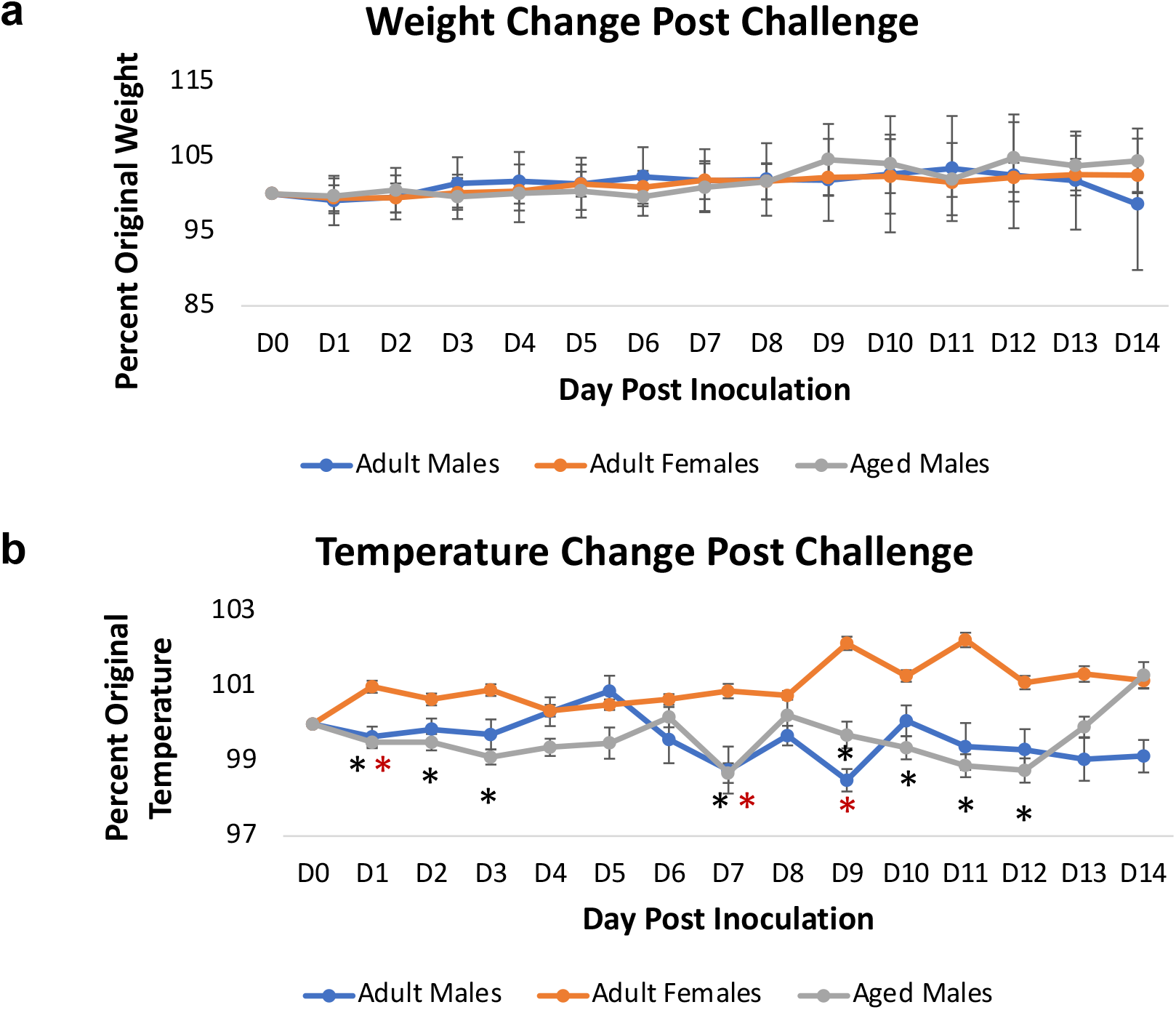
Intranasal SARS-CoV-2 infection in sexually intact ferrets leads to significantly lower temperature in adult males and aged males compared to adult females. **a** Sexually intact adult female, adult male, and aged male ferrets were intranasally inoculated with the Severe Acute Respiratory Syndrome coronavirus 2 (SARS-CoV-2) (10^6^ TCID_50_) and weight was recorded for 14 days post inoculation. **b** Temperature was also analysed day for 14 days after inoculation. Results show the mean of at least 6 ferrets per group. * indicates a p-value less than 0.05 determined by ANOVA comparing adult females to adult males or aged males. Error bars indicate +/- SD (weight) or SE (temperature).

### SARS-CoV-2 infection in the nasal turbinates is more severe and longer lasting in aged male ferrets

Infectious viral loads in COVID-19 patients have been correlated with increased disease sequelae and mortality ^53^. It has been shown that males and seniors have increased SARS-CoV-2 viral burden compared to younger females ^54,55^. Although these reports suggest males have higher viral burden, modeling viral dynamics in human cohorts is difficult due to the requirement for frequent sample acquisition to obtain an accurate picture of viral kinetics, including duration and the peak of replication. Therefore, we assessed the viral burden in the upper respiratory tract nasal washes to determine the temporal dynamics of viral load and shedding. We collected nasal wash samples as described and assessed viral shedding by both vRNA by qRT-PCR and infectious virus by TCID_50_ assays. Marked differences were observed among the groups. Infectious virus was present in the nasal washes of female ferrets on day 2 pi though vRNA persisted until day 7 pi (Fig. 2A). The vRNA values of female nasal washes ranged from 3-4 TCID_50_/mL (Log10) which was significantly lower compared to those of adult males and aged males that had maximum vRNA >7 TCID_50_/mL (Log10) at their respective apex. Conversely, adult males and aged males shed infectious virus until days 5 and 7, respectively. Nasal turbinates were then assessed as a possible virus source (Fig. 2B). vRNA and infectious virus were found in the nasal turbinates of female ferrets on day 2 pi alone, similar to titers seen in the adult and aged males. Although females were only positive on day 2 pi, adult and aged males were positive for virus at later time points. By day 7, adult males were positive for vRNA alone, whereas the aged males were positive for both vRNA and infectious virus in the nasal tissues. To determine if the antibody response was differentially affected by sex and age, we conducted microneutralization assays. Plasma collected from adult females (day 14 and 21 pi), adult males (day 14 pi only), aged males (day 14 and 21 pi) were evaluated. No differences were noted among the experimental groups. Ferrets were able to elicit SARS-CoV-2 neutralizing antibodies to average titres between 1:20 to 1:40 by day 14. All animals had detectable neutralizing antibodies except for one aged male (day 14) and one adult female (day 21) (Fig. 2C).

**Fig. 2:**
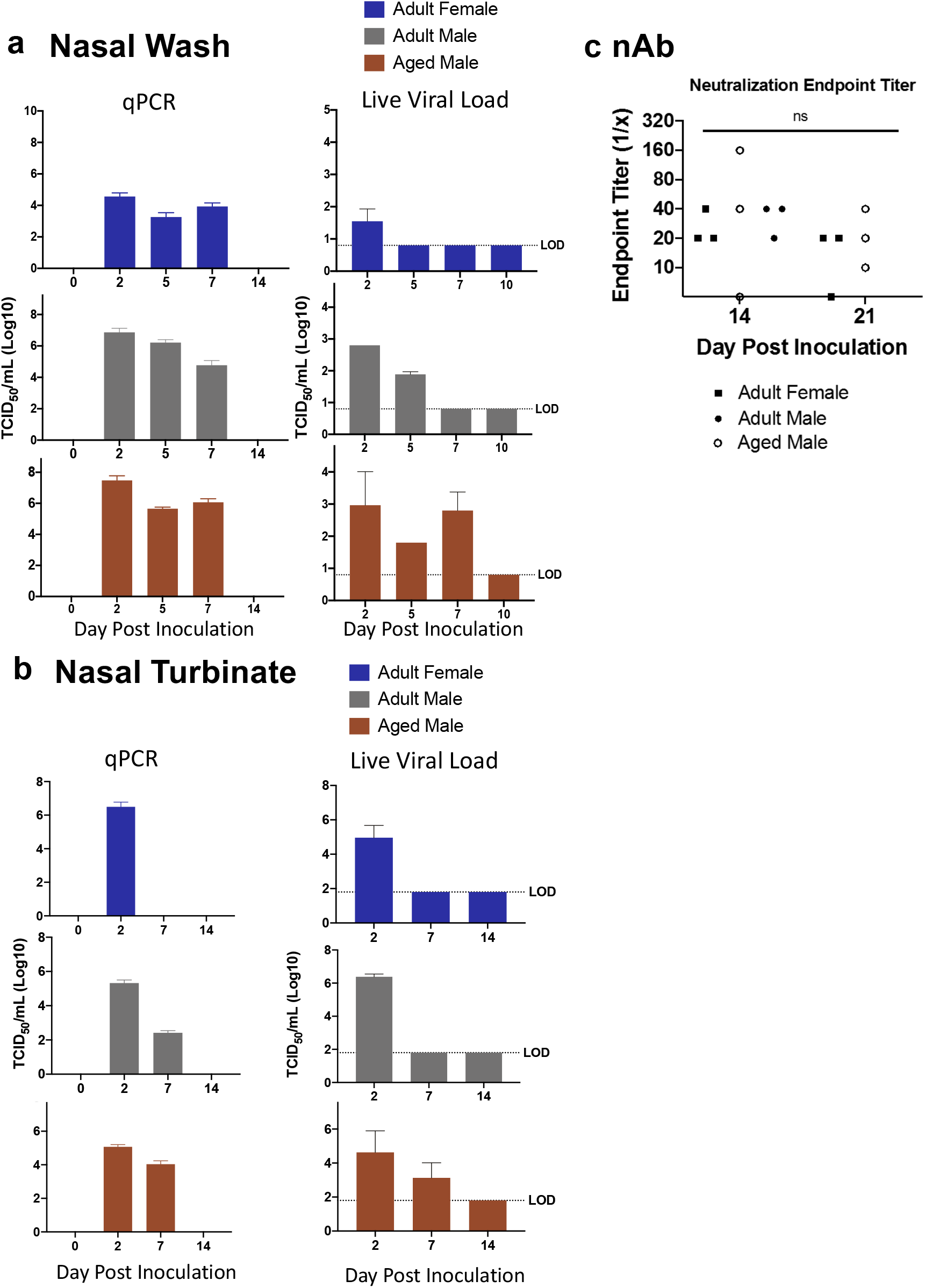
Prolonged SARS-CoV-2 viral presence occurs after infection in upper respiratory tract of aged male ferrets. **a** Nasal washes were collected were harvested to quantify SARS-Co-V-2 by qPCR and live viral load assay. Infectious titer of TCID_50_/mL was calculated using the Reed and Muench method. **b** vRNA and live virus was also quantified in the nasal turbinates. **c** Virus neutralization titers were determined by standard neutralization assays using plasma collected from all three groups on days 14 and 21 pi. LOD = limit of detection. Error bars represent standard error. N is 3 for all groups.

We assessed vRNA and infectious virus presence in tissues outside of the upper respiratory tract to determine if virus tropism was expanded outside the upper respiratory tract. Following infection, we collected the salivary glands, trachea, lung lobes (right cranial, right middle, right caudal, left cranial, left caudal, and accessory lobes), mediastinal lymph nodes, heart, kidney, liver, spleen, and large intestine (colon) from all three experimental groups on days 2, 7, and 14 pi. Viral presence by qRT-PCR showed adult females were positive for vRNA in the trachea, accessory lobe, and the colon on day 2 pi, whereas vRNA was found only in the colon (Large Intestine (LI)) of the adult males (Supplementary Figure 1). In contrast, vRNA was positive in all lung lobes as well as the LI in aged males (Supplementary Figure 1, bottom left panel). By day 7 pi, adult male and female animals had additional vRNA positive tissues; the number of vRNA positive tissues decreased at this time point in the aged males. All tissues were negative across all groups on days 14 and 21 (results not shown). All tissues positive for vRNA were then assessed for the presence of live virus by TCID_50_ titration assay. Although several tissues were positive for high quantities of vRNA, no tissues were positive for live virus. Taken together, these results show that males have a longer duration of viral replication and shedding from the upper respiratory tract, possibly affecting the lung lobes as well, which was also associated with age. These results suggest that females are more effective at limiting viral loads and replication over time.

Due to the differences we observed across the ferret groups in both vRNA and infections in the upper respiratory tract, we assessed infection-mediated pathology within nasal turbinate tissues (Fig. 3). Nasal turbinates were collected at necropsy from SARS-CoV-2-infected animals and were formalin-fixed, paraffin embedded, and H&E prepped (Fig. 3) or stained for viral antigen by IHC (Fig. 4). Tissues were visualized and images captured with the Leica DMI100 Brightfield microscope and camera. Nasal turbinates from non-infected animals showed ciliated epithelial cells with minimal infiltration of immune cells or evidence of tissue erosion (Fig. 3A). Moreover, control tissues from uninfected animals were negative for viral antigen staining (Fig. 4A).

**Fig. 3:**
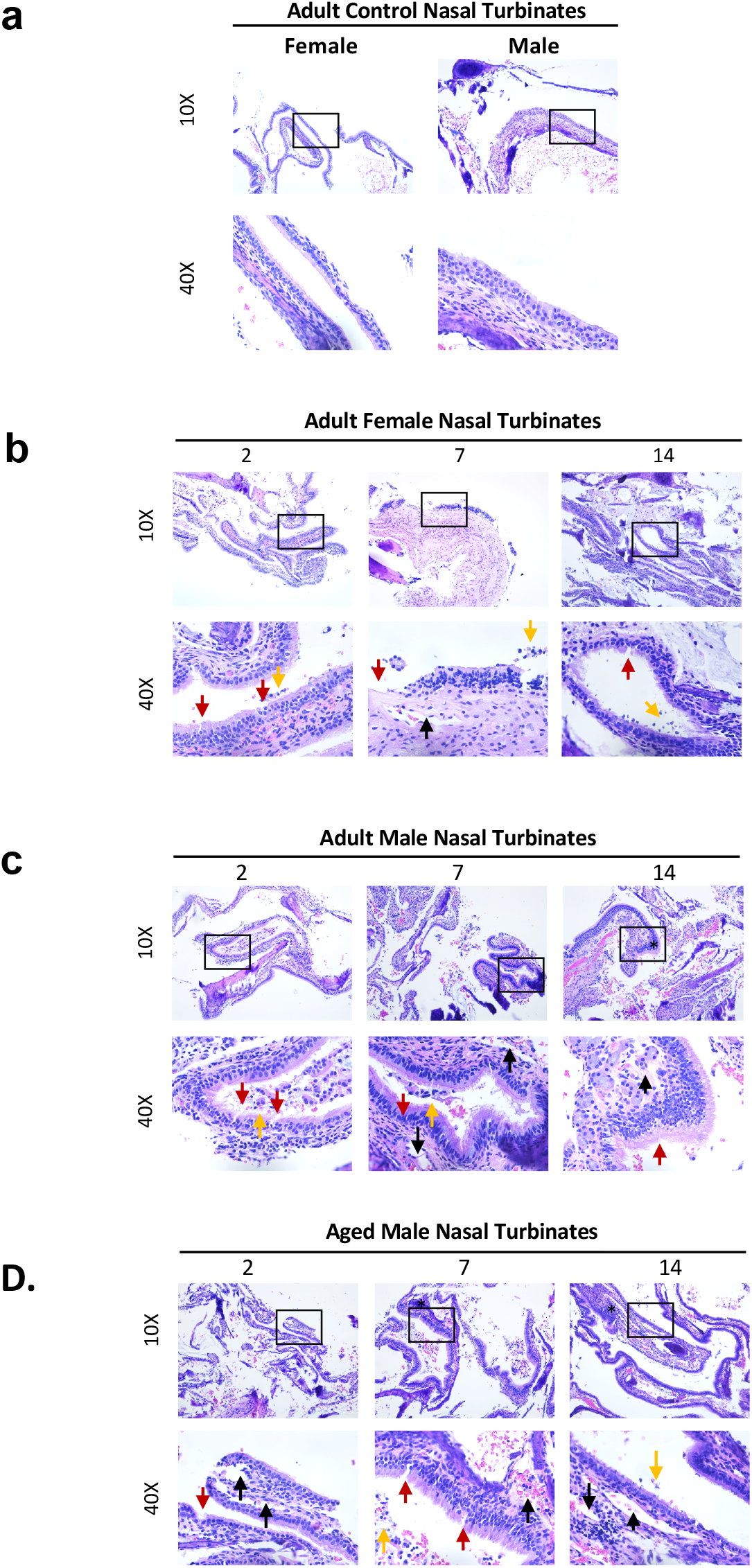
Aged male ferrets inoculated with SARS-CoV-2 have increased damage in the nasal turbinates compared to adult female and adult male ferrets. **a** Nasal turbinates were examined by H&E collected from uninfected control adult ferrets. **b** Histopathology of H&E stained nasal turbinates were also analysed in infected adult femal. **c** Histopathology of H&E stained nasal turbinates from infected adult male. **d** Histopathology of H&E stained nasal turbinates from infected aged male ferrets. The collected nasal turbinates were formalin-fixed and paraffin block embedded prior to slide mounting and staining by H&E. Stained tissues were visualized and images captured via the Leica DMI100 Brightfield microscope and camera. Images were captured at 10X and 40X. Images shown are representative of 3 animals per group, per timepoint. Black arrows indicate cavity formation; red arrows denote epithelial erosion; and orange arrows point to epithelial cell sloughing. Lines refer to loss of ciliated cells.

**Fig. 4:**
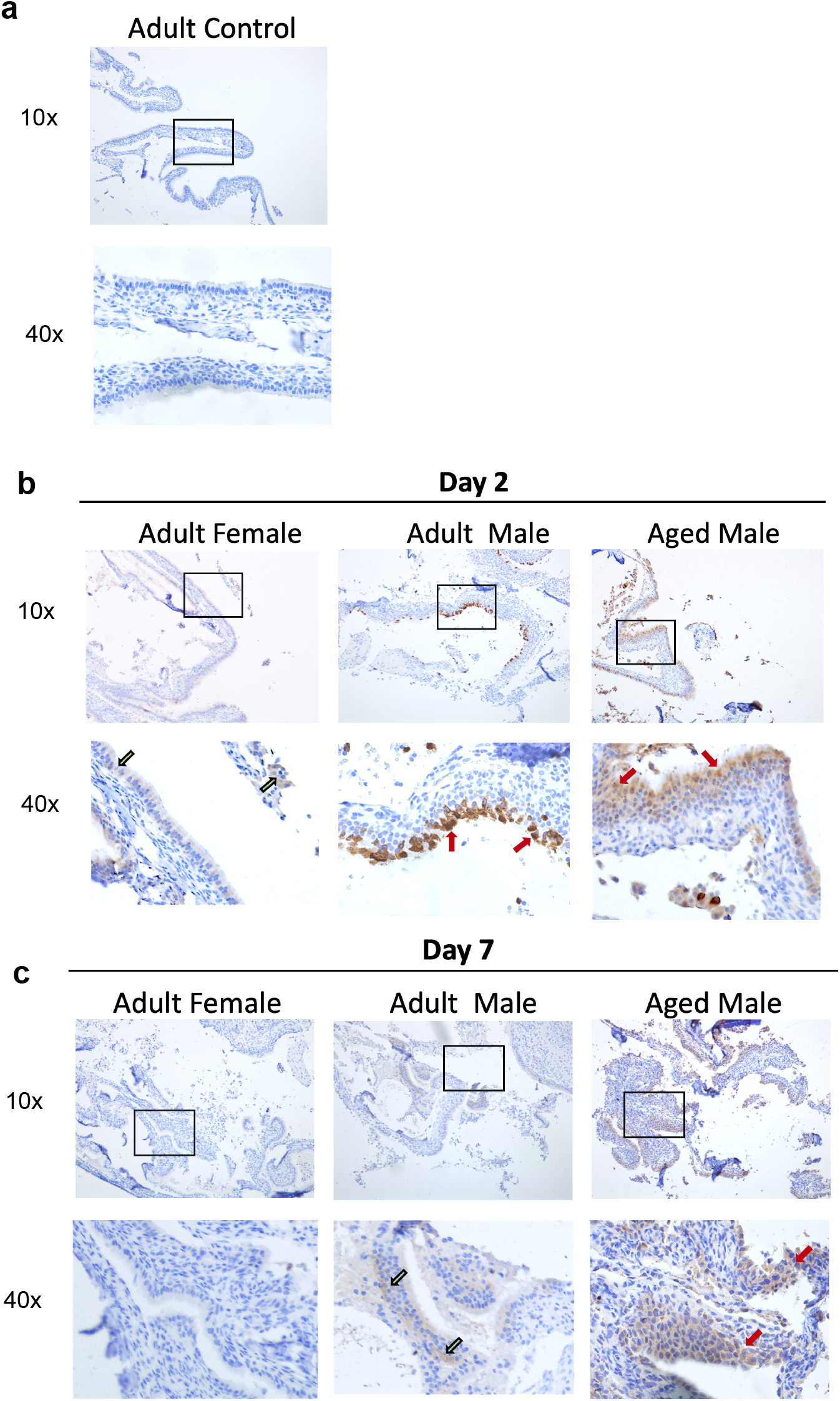
Aged male ferrets have increased and prolonged SARS-CoV-2 viral antigen presence in nasal turbinates. **a** Formalin-fixed nasal turbinates collected from uninfected ferrets were stained for SAARS-CoV-2 spike protein. **b** Nasal turbinates collected from adult female, adult male, and aged male ferrets on day 2 post SARS-CoV-2 inoculation. **c** Nasal turbinates collected from adult female, adult male, and aged male ferrets on day 7 post SARS-CoV-2 inoculation. Red arrows indicate high amounts of viral antigen staining and green arrows indicate low amount of viral antigen staining) by immunohistochemistry (IHC) using an anti-spike rabbit monoclonal antibody. Stained tissues were visualized and images captured using a Leica DMI100 Brightfield microscope and camera. Images were captured at 10X and 40X. Images shown are representative of 3 animals per group, per timepoint.

Turbinates from adult female ferrets had minimal immune cell infiltration and tissue architecture destruction day 2 pi (Fig. 3B). Ciliated epithelial cells were visible at this time point with minimal cilia loss. Mononuclear cell infiltration was first detected on day 7 and was most pronounced on day 14 pi as indicated by the asterisk in the females. Rhinitis, inflammatory cell infiltration, and loss of ciliated cell layers are evident at this time. Granulocytes can also be seen from days 7-14 pi. Adult females had minimal viral antigen staining in the pseudostratified columnar epithelium which was only observed on day 2 pi (Fig. 4B, left lower panel, indicated by green arrows).

Adult male ferrets had increased ciliated epithelial cell loss starting on day 2 pi (indicated by red arrows) compared to females (Fig. 3C). Cavity formation was first noted on day 7 (black arow) with concomitant edema and presence of immune exudate, monocytic cell infiltrate, disruption of cilia and cavitation, and heavy sloughing. Significant mononuclear cell infiltration, as well as signs of necrosis, was evident by day 14 pi as indicated by the asterisk (Fig. 3C, far right panels). Adult males had a high concentration of positively stained cells in the outer layer of the nasal olfactory epithelium on day 2 (Fig. 4B, middle panels, indicated by red arrows) which diminished by day 7 (Fig. 4C, green arrows).

Nasal turbinates from aged males had the most significant epithelial destruction and immune cell infiltration in all groups (Fig. 3D). Evidence for mononuclear cell infiltration was identified as early as day 2 pi (asterisk) and remained visible throughout the time course. Significant loss of ciliated cells was observed on day 2 pi (black line) with signs of erosion in the surface of the epithelial cells (red arrow), as well as cavity formation (black arrow), necrosis, and immune infiltration. Erosion could be seen on days 2 and 7 pi whereas cavity formation in the submucosa was evident on days 7 and 14. Exocytosis of mononuclear cells was consistently observed post infection. Orange arrows point to epithelial cell sloughing. Aged males also had high amounts of viral antigen in the nasal olfactory epithelium, but the staining was not limited to the outer layer (Fig. 4B, lower right panel, indicated by red arrow). Staining throughout the nasal olfactory epithelium was seen in aged males on day 7, which again reached the basement membrane (Fig. 4C, lower right panel, red arrow).

Together, all infected ferret groups showed damage to the nasal turbinates over the course of infection with evidence of viral antigen within the tissue. Adult males and aged males had early signs of mononuclear cell infiltration and cavity formation with damage to the tissue throughout the time course. Tissue analysis of viral antigen revealed more extensive, and a longer duration of, antigen in the aged males which was consistent with our viral load findings.

### Antiviral and Interferon Stimulated Gene upregulation in the nasal turbinates positive for SARS-CoV-2 were delayed in aged male ferrets but neurosensory and inflammatory signalling genes were increased at later time points

To identify candidate mechanisms underlying prolonged viral replication in SARS-CoV-2-infected aged male ferrets, we performed unbiassed, whole transcriptome transcriptome analysis. RNA was isolated from upper (nasal turbinates) and lower (right cranial and caudal lung lobes) respiratory tracts of control animals, infected adult female ferrets and aged male ferrets at days 2, 7, 14, and 21 following inoculation. Gene enrichment analysis plots, leveraging the Hallmark Signalling Pathway collection^56,57^ indicated significant differences in the longitudinal profiles of several prominent signalling pathways between SARS-CoV-2 infected adult females and aged males (Fig. 5). Immune-relevant signalling pathways that were significantly regulated for at least one time point compared to baseline for adult females or aged males included interferon-alpha, interferon-gamma, unfolded protein response, MTORC, MYC targets V2, DNA repair, PI3K AKT MTOR signalling, reactive oxigen species, IL2 STAT5 signalling, allograft rejection, apoptosis, inflammatory response, peroxisome, oxidative phosphorylation, IL6 JAK STAT3 signalling, and complement. Adult female and aged male ferrets had obvious differences in the longitudinal trends of several of these signalling pathways. Most noted were the early increase in interferon-alpha (IFN-a) and interferon-gamma (IFN-g) for the adult females on day 2 pi (Fig. 5A) whereas the aged males had no expression of interferons on day 2 which peaked later on day 7 pi (Fig. 5B). Also of note were the decreasing trends of inflammatory response associated pathways in the adult females on day 21 of reactive oxygen species, IL2 STAT5 signalling, complement, inflammatory responses, allograft rejection, and apoptosis. Conversely, aged males had statistical increases in pathways involved with inflammatory responses including oxdative phosphorylation, fatty acid metabolism, peroxisome, reactive oxygen species, MTORC and unfolded protein response. Once we had gained a perspective of the overall pathway level changes over time, we looked more closely at the top 50 DEGs by p-value (all p<0.05) in isolation by day post-infection to identify the individual genes represented in each pathway (Fig. 6). Interferon response and antiviral response genes most highly expressed in the female day 2 nasal turbinates included OASL, MX1, DHX58, RSAD2, ISG15, and CTSG. By day 7 pi, aged male ferrets had a shift in gene profiles with significant increases in CXCL9, CXCL10, CXCL11, OASL, RSAD2, IRF7, ISG15, ISG20, CAMP, and CTSG. Similar trends were also seen in analysis of the right cranial (Supplementary Figure 2) and right caudal (Supplementary Figure 3) lung lobes. Specifically, IFNG, IFI6, IRF7, DHX58, RSAD2, IFIT1, CMPK2, MX1, and USP18 were highly upregulated in the right cranial lung lobes of the females but not in males on day 2 pi. This was consistently observed in the right caudal tissue of females as well (RNF7, DHX58, IFI6, RSAD2, IFIT1, IFIT3, ISG15, and MX1). Select genes validated by qRT-PCR confirmed the trend of delayed interferon responses in aged males compared to adult females for both the upper and lower respiratory tract (Supplementary Figure 4). In addition, adult males were also investigated for regulation of interferon responses by qRT-PCR (Supplementary Figure 4). This analysis showed adult males have a unique signature of host responses (interferon and inflammatory gene expression) following SARS-CoV-2 infection. Specifically, IFN-a and IFN-g were regulated to similar expression levels as adult female ferrets but inflammatory genes such as IL-6 were regulated at intermediate levels. Taken together, these results suggest that together age and sex in males may bias the IFN responses to be delayed during SARS-CoV-2 infection in the respiratory tract.

**Fig. 5:**
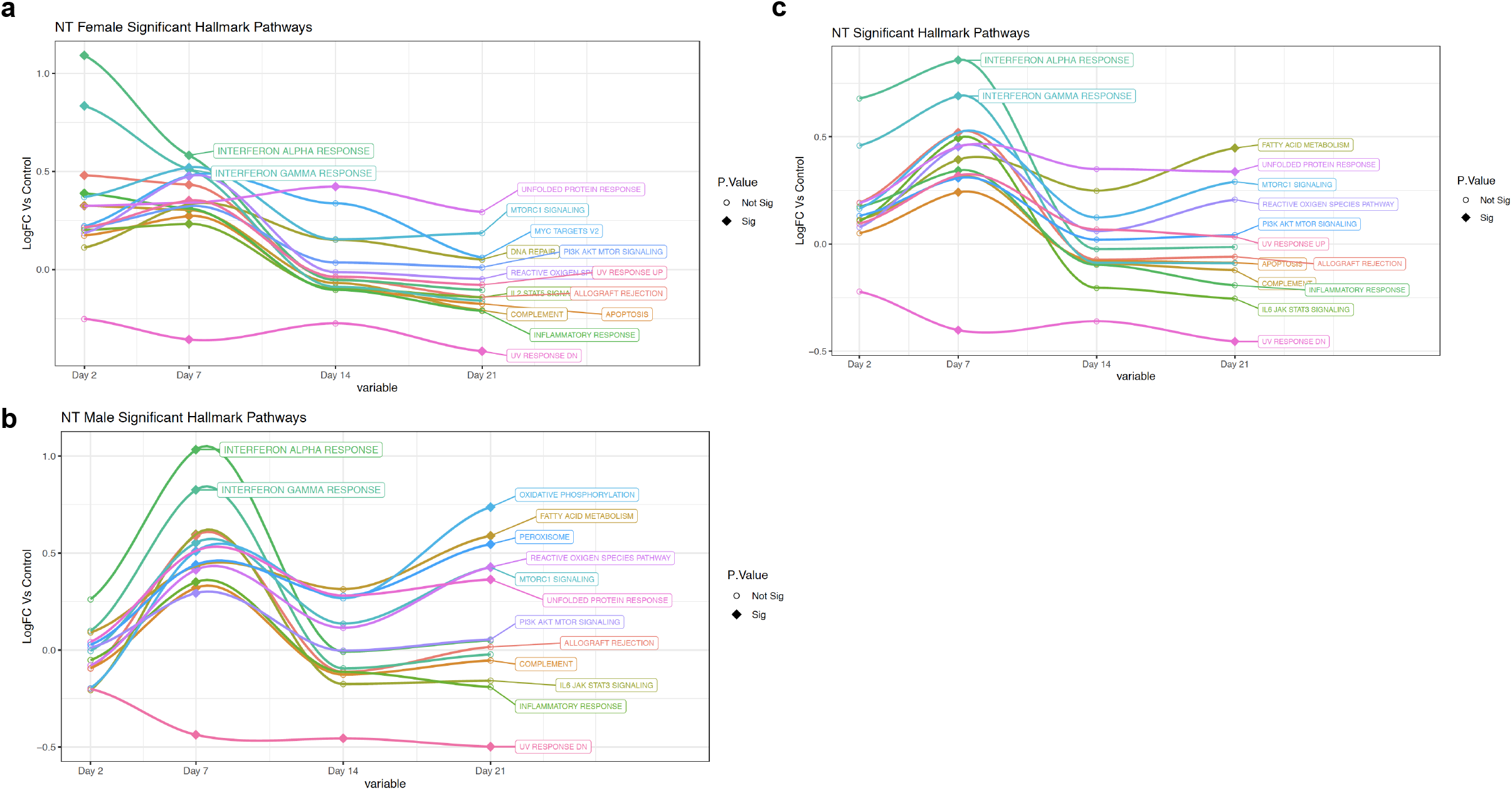
Unique signalling pathway regulation for aged males and adult females in the upper respiratory tract of ferrets infected with SARS-CoV-2. Longitudinal enrichment analysis was plotted following RNAseq of aged male and adult female RNA isolated from the nasal turbinates collected on day 0, 2, 7, 14, and 21 post SARS-CoV-2 inoculation. **a** Enrichment profiles were created using Hallmark Pathways for adult females. **b** The Hallmark enrichment profiles for aged males. **c** The Hallmark enrichment profiles for aagedd males. Signalling pathways which were statistically changed (FDR < 0.10) compared to baseline for at least one time point were included: Interferon-alpha, Interferon-gamma, Unfolded Protein Response, MTORC, MYC Targets V2, DNA Repair, PI3K AKT MTOR Signalling, Reactive Oxigen Species, UV Response, IL2 STAT5 Signalling, Allograft Rejection, Apoptosis, Inflammatory Response, UV Response DN, Peroxisome, Oxidative Phosphorylation, IL6 JAK STAT3 Signalling, and Complement. Filled diamonds represent a time point statistically changed compared to baseline. Open circles represents time points that are not statistically significant compared to baseline. Three ferrets per group per time point were analyzed with the exception of a poor sample in the adult female on day 7 which did not pass quality control.

**Fig. 6:**
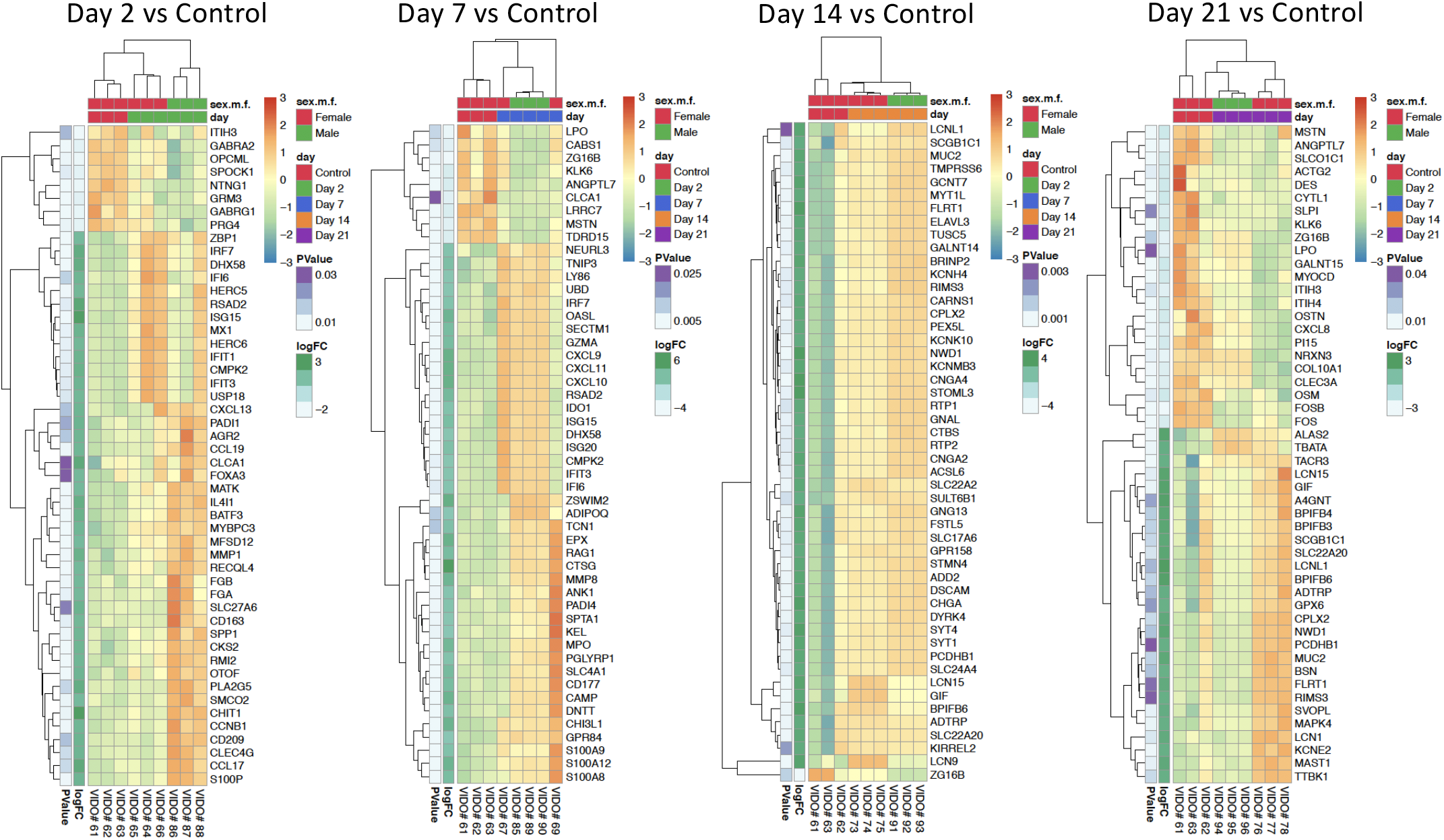
SARS-CoV-2 infected adult female ferrets have early upregulation of interferon responses in the nasal turbinates not observed in aged males while aged male ferrets have increased olfactory-associated genes at later time points. RNA extracted from infected aged male and adult female nasal turbinates were subjected to RNA sequencing on the Illumina platform to determine the host transcriptomic responses during SARS-CoV-2 infection. Hierarchical clustering of the Top 50 differentially expressed genes are respresented from day 2, 7, 14, and 21 post inoculation.

To further elucidate specific genes involved in the regulation of overrepresented pathways in nasal turbinate samples at the later time points, we examined specific genes regulated in the top 50 DEGs of nasal turbinate samples from days 14 and 21 pi. Interestingly, genes associated with neurological signalling and the olfactory response were overrepresented in the aged male ferrets on day 14 pi (Fig. 6). Twenty of the top 50 DEGs were associated with these pathways and included GNG13, SLC17A6, FSTL5, BPIFB6, MYT1L, FLRT1, BRINP2, RTP1, STMN4, DSCAM, CHGA, DYRK4, PCDHB1, SYT4, SYT1, CNGA2, ACSL6, STOML3, CARNS1, CPLX2, CNGA4, SLC22A20, and KCNMB3. Of note, GNG13 (taste recognition), SLC22A20 (odorant transporter), RTP1 (olfactory receptor binding), CNGA2 (associated with anosmia), and CNGA4 (transduction of odorant signals) have all been implicated in the olfactory responses. On day 21, neurological signalling-associated genes were still present, but expression profiles varied and were not sex specific. These results suggest that there is a sex- and age-biased recovery following SARS-CoV-2 infection in the upper respiratory tract that may affect sensory perception.

## Discussion

Here we investigated the influence of age and sex on clinical outcomes and host responses after SARS-CoV-2 infection using the ferret model. Importantly, by using ferrets with intact sex organs, we showed that aged male ferrets had lower temperature compared to baseline and had prolonged SARS-CoV-2 viral shedding after inoculation. Host transcriptome analysis suggested that aged males had delayed expression of antiviral genes such as MX1, IRF7, and ISG15 compared to adult females, which may have been responsible for the sex-dependent delay in viral clearance. Aged male ferrets also had increased expression of genes associated with odor detection at later time points, possibly indicative of key drivers of COVID-19 anosmia. COVID-19 caused by SARS-CoV-2 has significantly impacted both the male sex and older individuals. Our findings may inform human studies and suggest investigation of the interrelationship of viral shedding, interferon signalling, and olfactory responses in COVID-19 patients which may be biased in males and the elderly. The identification of prolonged viral shedding in older males may have implications in public health responses and the analysis of transmission events.

To study the contribution of sex and age to the development of severe COVID-19, we were able to parse the variables of age and sex to stratify clinical, virological, and immune outcomes post SARS-CoV-2 infection by utilizing ferrets with intact sex organs. Based on human clinical and epidemiological findings disaggregated by sex and age, we hypothesized that older male ferrets would develop severe disease post SARS-CoV-2 infection ^48–50,58^. Since older males are at higher risk of hospitalization and death from SARS-CoV-2 infection, we anticipated the establishment of a preclinical model with a severe COVID-19 phenotype that could be employed for vaccine and therapeutic development. In contrast, we found that none of our older or sex organ intact ferrets developed significant weight loss after SARS-CoV-2 infection, but temperature was significantly affected. We found that females developed a sustained mild fever similar to that reported for ferrets (possibly spayed and neutered) infected with SARS-CoV-2 in other studies ^59^. Conversely, all males in our study experienced a significant decrease in temperature with older males having statistically lower temperature than females compared to the younger males. Interestingly, temperature has been shown to be dysregulated in older patients infected with other respiratory viruses. As older patients do not consistently develop conventional temperature increases during infection of viruses such as influenza virus, elderly diagnoses have been missed, which eventually affects patient treatment and outcome ^60^. Our results suggest older male COVID-19 patients may also fail to be identified in early stages because of atypical clinical pictures. Currently, there are limited clinical definitions of COVID-19 per age group and sex. Defining the clinical picture for specific cohorts will be essential for proper diagnosis, treatment plans, and patient outcome.

Although SARS-CoV-2 continues to circulate globally, little is understood regarding host-specific viral dynamics and transmission. In our preclinical study, we found that there was a sex and age influence on SARS-CoV-2 in the upper respiratory tract. Male ferrets shed live SARS-CoV-2 virus longer than females and shedding virus from males increased by age. Interestingly, analysis of vRNA and live virus in the nasal turbinates day 2 pi did not indicate a quantitative difference of viral burden by sex or age, indicating that females and males were equally susceptible to infection and viral replication. Therefore, the differences that were observed among the groups in respect to virus durability and prolonged shedding conversely suggested there was a sex and age bias in the ability to control viral infection over time. The viral loads we detected in aged male ferrets were similar to those reported by other SARS-CoV-2 ferret studies (viral clearance ~ day 8 pi), leaving the possibility that the presence of female sex hormones or sex organs in our sexually intact female ferrets influenced the efficiency of viral clearance ^34^. In respect to human COVID-19, our results seemed to recapitulate human SARS-CoV-2 viral load demographics where older people as well as males have been tested to be SARS-CoV-2 positive for longer periods of time ^54,55^. Mouse models of SARS-CoV-1 as well as SARS-CoV-2 also indicated that viral burden in the respiratory tract of males and older animals is also significantly increased ^40,61,62^ suggesting similar influence of age and sex for SARS-like coronaviruses. In contrast to these findings, a preclinical study investigating the effect of age on SARS-CoV-2 morbidity using the Syrian hamster model found no difference in viral shedding of older and younger animals ^63^. This report may indicate that although the Syrian hamster model recapitulates lung disease associated with human COVID-19^38^, ferrets and mice may be a superior model for age-based studies.

We found an association between altered immune responses with prolonged viral load in aged males. The human COVID-19 demographics clearly indicate that older males are at higher risk of severe COVID-19 but this data does not give insight into the factors influencing disease. In respect to sex biases, biological and behavioural factors may both influence disease demographics ^64^. Our results leveraging a controlled preclinical animal model that removed behavioural factors suggested that biological sex strongly affected the outcome of SARS-CoV-2 infection. Recently, humoral responses as well as interferon signalling have been implicated in the sex biases of viral infection. Data from severe COVID-19 patients indicated that males and the elderly had increased neutralizing antibodies toward SARS-CoV-2, possibly suggesting a pathogenic humoral involvement or mechanism affecting antibody production during infection ^50^. Several other human and animal studies have reported that perturbed interferon responses are associated with SARS-CoV-2 infection, suggesting that dysregulated interferon signalling is a common mechanism driving disease ^58,65 66^. Although we did not find statistical differences in neutralizing antibody titers between males and females or by age, we did observe a significant delay in interferon responses in aged male ferrets. Specifically, adult female ferrets had earlier antiviral responses characterized by increases in OASL, MIX1, DHX58, RSAD2, ISG15, and CTSG, which would be essential for viral clearance. These genes were absent in the aged males at day 2 pi. In agreement with our findings, one study has shown that male COVID-19 patients had a skewed interferon response ^58^. This report by Bastard and colleagues showed a significant number of autoantibodies elicited toward type I interferons, possibly leading to interferon dysregulation. Previous studies have also suggested a role for Toll-like Receptor 7 (TLR7) in sex-biased interferon signalling during viral infections ^58^. TLR7, a viral nucleic acid pathogen recognition receptor, is located on a region of the X chromosome which is known to have a lower level of chromosomal inactivation, allowing females to have stronger interferon responses during viral infection due to gene dosing ^49,67–69^. Our data suggested that male ferrets had an atypical response to SARS-CoV-2 which was clinically characterized by a period of hypothermia and molecularly characterized with delayed interferon gene upregulation. Taken together, these reports may represent mechanisms contributing to the prolonged viral presence and subsequent dysregulated interferon responses we found in the upper respiratory tract of older males. Future studies should explore the sex-specific regulation of interferon responses with regard to the involvement of type I interferon-specific antibodies or TLR7 signalling in viral replication and clinical morbidity ^58,68,70^.

Anosmia, or loss of smell, has been reported as one of the more common symptoms of COVID-19 ^71,72^. Studies have found that between 33.9% and 68% of COVID-19 patients experienced symptoms associated with loss of smell ^71–73^. Sex biases have also been reported for COVID-19 associated anosmia with as many as 72% of COVID-19 anosmia patients being women ^72,73^. Considering that males are disproportionately affected with severe COVID-19 after SARS-CoV-2, it is intriguing that women have increased representation of anosmia. Interestingly, our gene expression analysis of SARS-CoV-2 infected nasal turbinates indicated that males had significant increases in genes associated with smell and sensory perception following virus resolution which were not present in female profiles. Since we did not measure olfactory responses in the infected ferrets, we are unable to conclude if loss of smell occurred and if there was an associated sex bias. Although measurement of smell was outside the scope of our study, our transcriptome analysis suggested that odorant responses were affected post SARS-CoV-2 infection and in a sex-biased manner. Specifically, the genes GNG13 (taste recognition), SLC22A20 (odorant transporter), RTP1 (olfactory receptor binding), CNGA2 (associated with anosmia), and CNGA4 (transduction of odorant signals) were differentially regulated in males. As males have a lower incidence in anosmia, it is possible that these genes are associated with smell retention during infection, but this hypothesis should be investigated further. Little is understood regarding the molecular aspects of olfactory dysfunction and in particular the cause of COVID-19 related anosmia but these profiles may give insight into anosmia regulation or protection.

Our study was limited by lack of an aged female ferret group, due to lack of a supply of these animals early in the pandemic. Therefore, our conclusions focus on the suggestion that early antiviral intervention may be more beneficial for males compared to females infected with SARS-CoV-2. Moreover, aged male ferrets had higher viral load that persisted longer in their upper respiratory tracts and that in turn was associated with uniquely delayed antiviral gene expression. While further research is needed to confirm the target genes we have identified, our study represents a unique transcriptomic resource and established animal model for futher research.

There is a critical need to understand the mechanisms underlying severe COVID-19 and how host specific factors play a role in disease progression and outcome. Here we found that aged male ferrets had higher viral load that persisted longer in their upper respiratory tracts, which was associated with delayed antiviral gene expression. Taken together, these findings suggest that early antiviral intervention may be more beneficial for males compared to females infected with SARS-CoV-2. Further sex-aggregated analysis of antiviral use in clinical studies may be beneficial to understanding the outcome of antiviral treatment during COVID-19 for compounds such as remdesivir. More work is needed to expand on these conclusions and identify whether these mechanisms are also at play in humans and if older males may be a source of super spreader events due to prolonged viral shedding.

## Methods

### Ethics Statement

All work was conducted in accordance with the Canadian Council of Animal Care (CCAC) guidelines, AUP number 20200016 by the University Animal Care Committee (UACC) Animal Research Ethics Board (AREB) from the University of Saskatchewan. For ferret manipulation, 5% isoflurane anesthesia was used with all efforts to minimize suffering.

### Virus

The SARS-CoV-2 isolate /Canada/ON/VIDO-01-2020 used for infections and *in vitro* assays was isolated from a patient presenting at a Toronto hospital upon returning from Wuhan, China ^74^. The viral stock grown in vDMEM (DMEM (Dulbecco’s Modified Eagle Medium) (*Wisent Bioproducts (Cat # 319-005-CL)*), 2% fetal calf serum (*Wisent Bioproducts (Cat # 090-150)*), 5 mL 100x Penicillin (10,000 U/mL)/Streptomycin (10,000 μg/mL), and 2 μg/mL TPCK-trypsin) was from the second passage (GISAID – EPI_ISL_425177). All work with SARS-CoV-2 live virus was performed in a CL3 facility at VIDO-InterVac (Saskatoon, Saskatchewan, Canada).

### Animals, Infections, and Tissue Collection

Adult female (1 year), adult male (1 year) and aged male (2 years) ferrets with intact sex organs were purchased from Triple F Farms (Gillett, PA, USA). Ferrets were anesthetized for intranasal SARS-CoV-2 at 10^6^ TCID_50_ infections. On day 2, 7, 14, and 21 post infection, ferrets were terminally bled by intra-cardiac puncture and humanely euthanized. Following infection, weight, temperature, and clinical signs were monitored for 14-21 days. Weight and temperature were calculated as a percentage of original weights.

### Viral Titers

Nasal washes (1 ml) were collected post infection in anesthetized ferrets. Tissues collected at necropsy were homogenized in serum free DMEM using a Qiagen TissueLyzer. A 1:10 dilution series of sample was established in viral growth media (vDMEM) for TCID_50_ virus infection titration assays. Viral load was calculated by determining the 50% endpoint using the Reed-Muench method after observing cytopathic effect (CPE) on day 5 after cell inoculation.

### Viral Neutralization Assay

Plasma was heat-inactivated at 56°C for 30 m and then serially diluted 1:2. Virus was diluted to 100 TCID_50_ in vDMEM and used at a 1:1 ratio to plasma, incubated at 37°C for 1 h and added to cultured Vero-76 cells in 96 well plates for 1 h at 37°C. Inhibition of CPE was observed and recorded on day 5 post infection.

### RNA extraction and quantitative Real-Time PCR (qRT-PCR)

Tissue RNA was extracted using the Qiagen© RNeasy Mini kit cat. # 74106 (Qiagen, Toronto, Canada) according to the manufacturer’s instructions. vRNA was extracted from nasal washes using the Qiagen© QIAamp Viral RNA Mini Kit cat. # 52904 (Qiagen). All host qRT-PCR was performed in triplicate on cDNA synthesized as previously described ^44^ (Table 1). vRNA was quantified by Qiagen© Quanti-fast RT probe master mix (Qiagen, Toronto, Canada) using primer/probe sets specific for the SARS-CoV-2 E gene. The reactions were performed on a StepOnePlus™ Real-Time PCR System in a 96-well plate (Thermo Fisher) as previously described ^75^.

**Table 1.**
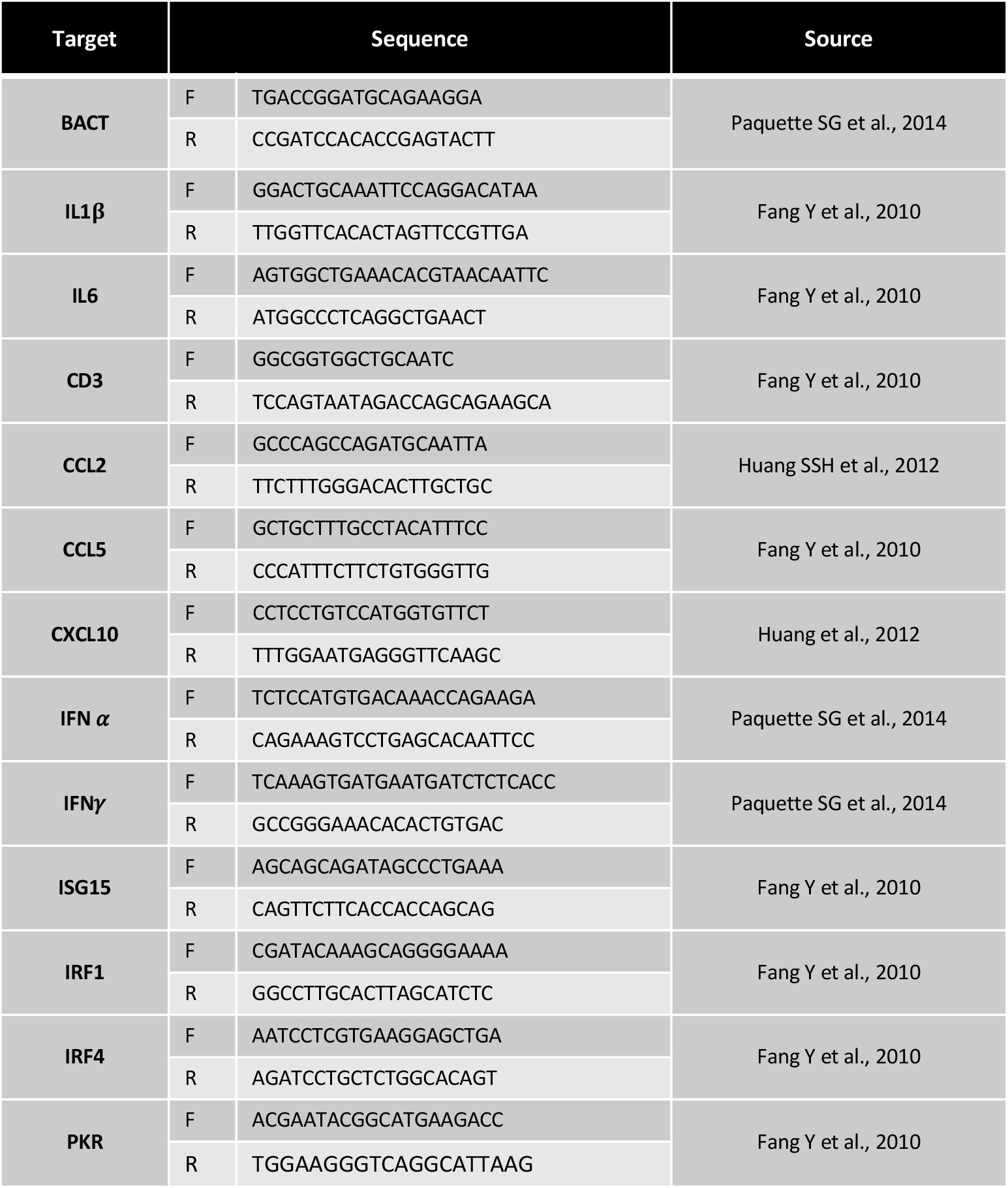
qRT-PCR primers used for host response validation.

### Histopathology and immunohistochemistry

The tissues collected for histopathology were submersed in formalin for +1 week. Formalin-fixed tissues were paraffin-embedded, sectioned, slide-mounted, and stained at Prairie Diagnostic Services (Saskatoon, Saskatchewan). Tissue samples were stained with hematoxylin and eosin or immunostained to detect SARS-CoV-2 antigen using VIDO-InterVac’s polyclonal rabbit anti-SARS-CoV-2 AS20-014 (1:400 dilution) and detected using an HRP-labelled polymer detection reagent (EnVision+ System – HRP Labelled Polymer, Agilent Technologies Canada Inc., Mississauga, ON).

### RNAseq

RNA was assessed with a Fragment Analyzer (Agilent) using the Standard Sensitivity RNA kit. Total RNA was normalized to 100 ng prior to random hexamer priming and libraries generated using the TruSeq Stranded Total RNA - Globin kit (Illumina, San Diego, USA). Libraries were assessed on the Fragment Analyzer using the High Sense Large Fragment kit and quantified using a Qubit 3.0 fluorometer (Life Technologies). Sequencing was performed on a NovaSeq 6000 (Thermo Fisher) using a 50 base-pair, paired-end run for a minimum of 30 million reads per sample.

### RNAseq Analysis

Raw demultiplexed fastq paired end read files were trimmed of adapters and filtered using the program skewer^76^ to remove any with an average phred quality score of less than 30 or a length of less than 36. Trimmed reads were aligned (HISAT2^77^) to the *Mustela putorius furo* NCBI reference genome assembly version MusPutFur1.0 and sorted using SAMtools. Reads were counted and assigned to gene meta-features using the program featureCounts ^78^ (Subread package). Count files were imported into the R programming language and assessed for quality control, normalized, and analyzed using the limma-trend method ^79^ for differential gene expression testing and the GSVA^80^ library for gene set variation analysis. Sequence and gene expression data are available at the Gene Expression Omnibus (accession number GSE160824).

### Statistical Analysis

Unpaired, unequal variance, two-tail Student’s *t*-test or one-way ANOVAs were conducted using GraphPad Prism8 (San Diego, USA). A *p* value of ≤ 0.05 was considered statistically significant.

## Data availability

Sequence and gene expression data are available at the Gene Expression Omnibus (accession number GSE160824).

## Acknowledgments

A. Kelvin is funded by the Canadian 2019 Novel Coronavirus (COVID-19) Rapid Research Funding initiative the Canadian Institutes of Health Research (CIHR) (grant numbers OV5-170349, VRI-172779, and OV2 – 170357) and Atlantic Genome/Genome Canada, Scotiabank COVID-19 IMPACT grant, and the Nova Scotia COVID-19 Health Research Coalition. D. Falzarano is funded by the Canadian Institutes of Health Research (CIHR), grant number OV5-170349. M. Cameron, C. Cameron, and B. Richardson are supported by NIH/NIAID (3R01AI129709-03S1) and the Nord Family Foundation, Amherst, Ohio. R. D. Pechous is supported by NIH/NIAID (award R56AI153252). J. Kindrachuk is funded by a Tier 2 Canada Research Chair in the Molecular Pathogenesis of Emerging and Re-Emerging Viruses provided by the Canadian Institutes of Health Research (Grant no. 950-231498). C. Richardson is funded by a CIHR COVID-19 Rapid Funding Opportunity VR1-172779. The authors would like to thank the tremendous efforts of the VIDO-InterVac veterinary staff including Dr. Colette Wheler. We also thank the staff of the CWRU Applied Functional Genomics Core for performing the RNA-Sequencing assays. Operational support at VIDO-InterVac is provided in part by the Canadian Foundation for Innovation through the Major Science Initiates Fund and by Innovation Saskatchewan.

This article is published with the permission of the Director of VIDO-InterVac.

## Author Information

### Contributions

Conceptualization: A. A. Kelvin; Investigation: A. A. Kelvin, M. E. Francis, J. Lew, D. Falzarano; Analysis: A. A. Kelvin, M. E. Francis, B. Richardson, R. D. Pechous, J. Kindrachuk; C. M. Cameron, M. J. Cameron, C. Richardson, M. Rioux; Writing: A. A. Kelvin, M. McNeil, M. Rioux, M. Foley, A. Ge, V. Gerdts

## Ethics declarations

Authors declare no competing interests.

