## Supplementary Data for "Male sex and age biases viral burden, viral shedding, and type 1 and 2 interferon responses during SARS-CoV-2 infection in ferrets"

**Supplemental Figure Legends**

**Supplementary Figure 1. Aged male ferrets show SARS-CoV-2 vRNA in respiratory tract and large intestine on day 2 post inoculation.** Adult female, adult male, and aged male ferrets were inoculated intranasally with $10^6$ TCID50 of SARS-CoV-2. Animals were removed from the study on day 0 prior to infection as well as days 2 and 7 post infection. Salivary gland (SG), trachea (T), right cranial lung (RCrL), right middle lung (RML), right caudal lung (RCaL), left cranial lung (LCrL), left caudal lung (LCaL), accessory lung (AL), mediastinal lymph node(MLN), heart (H), kidney (K), liver (L), spleen (S), and large intestine (LI) were harvested. Isolated RNA was assessed for the presence of the SARS-CoV-2 with the Qiagen Quanti-Fast RT probe master mix and primer/probe sets specific for SARS-CoV-2 E gene. An equivalent TCID50/mL was calculated based on CT values corresponding to a standard curve with virus of known titer. Error bars represent standard error. Three animals per group were analyzed for all groups.

**Supplementary Figure 2. Host response profiling analysis of the upper lung lobe (right cranial lung) in SARS-CoV-2 infected ferrets indicated that adult female ferrets have early activation of interferon stimulated genes not apparent in aged males.**

RNA isolated from the right cranial lung lobe of SARS-CoV-2 infected adult female and aged male ferrets was subjected to transcriptome analysis on the Illumina RNAseq platform. Samples were analyzed by day pi (control in *red*, day 2 in *green*, day 7 in *blue*, day 14 in *orange*, day 21 in *purple*) and sex (female in *red* and male in *green*) for the sample type of right cranial lung. Hierarchical clustering of the Top 50 DEGs indicated the induction of interferon response genes in the upper lung lobe of adult females although infectious virus was not detected in the lungs of any ferrets. Log fold change and p values were calculated and are evident on the left side of the clustergram for each gene of interest.

**Supplementary Figure 3. Host response profiling analysis of the lower lung lobe (right caudal lung) in SARS-CoV-2 infected ferrets indicated that adult female ferrets have early activation of interferon stimulated genes not apparent in aged males.**

RNA isolated from the right caudal lung lobe of SARS-CoV-2 infected adult female and aged male ferrets was subjected to transcriptome analysis on the Illumina RNAseq platform. Samples were analyzed by day pi (control in *red*, day 2 in *green*, day 7 in *blue*, day 14 in *orange*, day 21 in *purple*) and sex (female in *red* and male in *green*) for the sample type of right caudal lung. Hierarchical clustering of the Top 50 DEGs indicated the induction of interferon response genes in the lower lung lobe of adult females although infectious virus was not detected in the lungs of any infected ferrets. Log fold change and p values were calculated and are evident on the left side of the clustergram for each gene of interest.

**Supplementary Figure 4. Validation of transcriptome analysis by qRT-PCR indicated aged male ferrets have delayed CXCL10 and interferon response in the nasal turbinates and lung after SARS- CoV-2 infection.** qRT-PCR was performed on RNA extracted from nasal turbinate (A) and lung tissue (B) from SARS-CoV-2 inoculated adult female, adult male, and aged male ferrets. Samples were assessed for IL-1*b*, IL-6, CXCL10, CCL2, CCL5, IFN-*g*, CXCL10, PKR, CD3, IFN-*a*, IRF1, IRF4, and ISG15 with specific ferret primers (Table 1). Fold change was calculated via $\Delta\Delta$Ct against controls with BACT as the housekeeping gene. Error bars represent standard error. Three ferrets per group was used for the analysis of all time points. * represents a significant difference as determined by Student's t-test comparing against female as control.
